# Prodrug defiance reveals logic-based strategies for treating bacterial resistance

**DOI:** 10.1101/556951

**Authors:** Brandon Alexander Holt, Isabel Curro, Gabriel A. Kwong

## Abstract

Classifying the mechanisms of antibiotic failure has led to the development of new treatment strategies for killing bacteria. Among the currently described mechanisms – which include resistance, persistence and tolerance – we propose defiance as a subclass of antibiotic failure specific to prodrugs. Using locked antimicrobial peptides (AMP) that are activated by bacterial proteases as a prototypic prodrug, we observe that although treatment eliminates bacteria across the vast majority of environmental conditions (e.g., temperature, concentration of growth nutrients), bacteria spontaneously switch from susceptibility to defiance under conditions that alter the competing rates between bacterial proliferation and prodrug activation. To identify the determinants of this switch-like behavior, we model bacteria-prodrug dynamics as a multi-rate feedback system and identify a dimensionless quantity we call the Bacterial Advantage Heuristic (*BAH*) that perfectly classifies bacteria as either defiant or susceptible across a broad range of treatment conditions. By recognizing that the bacterial switch from susceptibility to defiance behaves analogously to electronic transistors, we construct prodrug logic gates (e.g., AND, OR, NOT, etc.) to allow assembly of an integrated 3-bit multi-prodrug circuit that kills defiant bacteria under all possible combinations of *BAH* values (i.e., 000, 001, …, 111) that represent a broad range of possible treatment conditions. Our study identifies a form of bacterial resistance specific to prodrugs that is described by a predictive dimensionless constant to reveal logic-based treatment strategies using multi-prodrug biological circuits.

## INTRODUCTION

The rise of multidrug-resistant bacteria coupled with the lack of newly developed antibiotics has created a serious public health threat ^1, 2^. Understanding the causes of antibiotic failure inspires the development of new drugs and informs clinical treatment strategies ^3–5^. These failure mechanisms are broadly classified into three distinct categories (i.e., resistance, persistence, and tolerance), which are characterized by the change in drug concentration and exposure time required to kill bacteria ^6^. For example, resistance is characterized by genetic mutations or phenotypic changes which result in bacteria requiring significantly higher concentrations of antibiotic (minimum inhibitory concentration, MIC) to be lethal. By contrast, bacteria exhibiting tolerance or persistence require increased drug exposure time (minimum duration for killing, MDK), with the latter exhibiting a biphasic killing profile (Fig. 1A). Further, bacteria populations become tolerant through environmental conditions or genetic mutations, whereas persistence is an actively maintained, non-heritable state exhibited by a subpopulation of bacteria. Discriminating between these survival strategies employed by bacteria is crucial for the development of new drugs and clinical treatment decisions ^6^.

**Figure 1.**
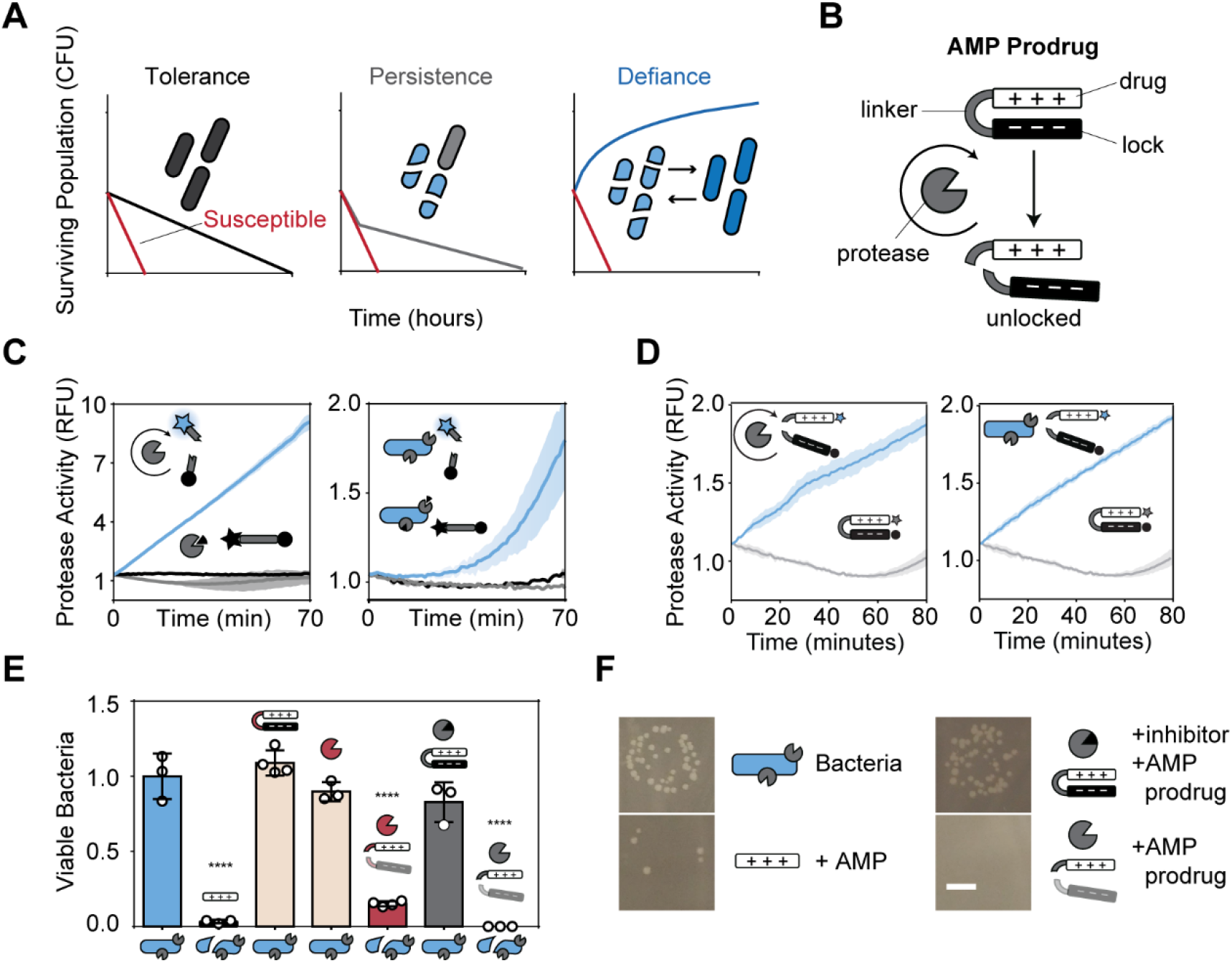
A bacteria-activatable prodrug targets *E. coli* protease OmpT. (**A**) Schematic comparing time-dependent killing profiles of bacteria exhibiting tolerance, persistence, or defiance. Red line represents killing profile of susceptible bacteria. (**B**) A cationic AMP drug (RRRRRRRRR, white rectangle) locked by an anionic peptide lock (EEEEEEEEEEEEE, black rectangle) with a protease-cleavable linker (RRS|RRV, grey u-shape) is activated by OmpT protease (grey pacman) activity. (**C**) Cleavage assay measuring the activity of recombinant OmpT (grey protease) (left) and OmpT expressed on the surface of E. coli (right) against fluorescence-quenched linker substrates (blue lines). Negative control samples contain the inhibitor aprotinin (black triangle, black lines) or linker substrates only (grey lines). Shaded regions represent standard deviation (n = 3). (**D**) Cleavage assay measuring the activity of recombinant OmpT (left) or OmpT expressed on the surface of E. coli (right) against fluorescently labeled hairpin prodrugs (blue lines) or hairpin prodrugs only control (grey lines). Shading represents standard deviation (n = 3). All cleavage assays plotted as fold change in RFU from initial time point. (**E**) Bacteria viability assay quantifying drug toxicity relative to untreated bacteria control (blue bar). Positive control for AMP toxicity (black bar). Negative control for locked AMP or TEV protease alone (tan bars). Positive control for TEV protease with locked AMP (substrate: ENLYFQ|G, specific to TEV protease) (red bar). Negative control for locked AMP (substrate: RRSRRV, specific to OmpT) with OmpT inhibitor, aprotinin (grey bar). Experimental condition of bacteria treated with locked AMP activated by natively expressed OmpT (white bar). All values normalized and statistically compared to bacteria only control. Error bars represent standard deviation (n = 3). (**F**) Representative images of bacterial plates used to quantify viability with schematic legend (scale bar = 4 mm). * < 0.05, ** < 0.01, *** < 0.001, and **** < 0.0001.

Antibiotic success is markedly improved by proper titration of drugs, as overdosing leads to off-target toxicity and underdosing increases the likelihood of pathogens developing resistance^7^. However, optimal drug doses are difficult to achieve over the course of treatment because infection burden changes dynamically over time, creating a moving target^7–8^. Prodrugs, which represent ∼10% of all FDA-approved drugs in the last decade, are a promising solution because they may be automatically titrated by a disease-related activation mechanism, increase bioavailability, and reduce the risk of off-target effects ^9^. For example, prodrugs such as ganciclovir^10^ and isoniazid^11^ are administered as biologically inactive forms and are enzymatically or chemically activated by the pathogen. Here, we use prototypic auto-titrating prodrugs comprising cationic antimicrobial peptides (AMP), which evolved as a part of innate immunity and display broad-spectrum and prokaryote-specific antimicrobial activity ^12^. Cationic AMPs act by disrupting bacterial membranes and inducing inflammatory responses ^*13*^, but suffer from off-target toxicity and low stability *in vivo* ^14, 15^. Our AMP prodrugs are activated directly by bacterial proteases, which are a major cause of virulence ^16–21^ during infections and important targets for antibiotic development ^22^. Of these proteases, outer membrane protease T (OmpT) is a membrane-bound bacterial protein that is widely conserved across gram-negative bacteria of the *Enterobacteriaceae* family and recognizes a variety of targets that contribute to its virulence (e.g., plasminogen)^18, 19, 23–25^. In this work, we design cationic AMPs that are locked by charge complexation with anionic peptides and connected with a protease-cleavable linker substrate ^26–28^ that is recognized by *E. coli* protease OmpT, such that increasing concentrations of bacteria activate higher concentrations of free drug.

While AMP prodrugs eliminate the majority of bacteria, we observe bacteria populations exhibiting a phenotype we named “defiant”, proliferating in the presence of active drug by consistently outpacing prodrug activation at all stages of growth (i.e., log phase, stationary phase). To understand the mechanism by which bacteria become resistant to AMP prodrugs, we built a system of coupled ordinary differential equations (ODE) that model the dynamics of the bacteria-prodrug competition. We use this model to identify a dimensionless quantity – the Bacterial Advantage Heuristic (*B.A.H.*) – which predicts with perfect accuracy whether bacteria will become defiant under a broad range of environmental conditions. At low *BAH* values, bacteria are susceptible to elimination, but beyond a critical *BAH* threshold, the bacteria switch to a state of defiance, in which the bacteria proliferate in the presence of activated drug. To combat prodrug failure due to defiance, we design biological circuits that eliminate bacteria under all conditions (*BAH* values) by recognizing that AMP prodrugs behave analogously to transistors: above a critical *BAH* threshold (i.e., gate voltage), bacteria (i.e., input current) pass through the prodrug and exhibit defiance. We use this framework to build logic gates with prodrug transistors, which we use to construct a multi-prodrug circuit that outputs 0 (i.e., dead bacteria) under all eight combinations of three input *BAH* values. These quantitative mechanistic insights will inform future drug design and treatment protocols to combat antibiotic resistance.

## RESULTS

### A bacteria-activatable prodrug targets *E. coli* protease OmpT

To construct bacteria-activated prodrugs, we synthesized cationic (polyarginine) AMPs in charge complexation with anionic peptide locks (polyglutamic acid) by a linker peptide (RRS|RRV) specific for the ubiquitous bacterial protease OmpT^26^. Upon proteolytic cleavage of the linker, the hairpin prodrug is unlocked to release free AMP, creating a mechanism for auto-titration (Fig. 1B). To demonstrate linker specificity for OmpT, we synthesized an activity probe^24, 29–33^ with free linker peptides containing a fluorophore-quencher pair and detected OmpT activity in samples incubated with recombinant OmpT or live *E. coli* culture. Conversely, we observed no activity in samples containing the serine protease inhibitor, Aprotinin, which inhibits OmpT when present in micromolar concentrations ^23^ (Fig. 1C). We observed similar cleavage activity using this linker substrate when fully integrated into hairpin AMP drug-lock complexes, confirming that linker presentation within a constrained conformational state did not affect cleavage activity by OmpT (Fig. 1D). To measure cytotoxicity of unlocked drug, we dosed bacteria with free AMP and observed significant reduction in colonies compared to untreated controls (blue bars) (Fig. 1E, F). To confirm prodrug specificity, we synthesized AMP drug-lock complexes using linker peptides specific for OmpT or Tobacco Etch Virus Protease (TEV), which exhibits orthogonal protease specificity ^34^. We observed elimination of bacteria only in samples containing OmpT-specific AMP prodrugs (grey bars) or samples treated with both TEV and TEV-specific AMP prodrugs (red bars). All control samples containing either TEV-specific prodrug alone or Aprotinin inhibitor did not significantly reduce bacteria load (Fig. 1E, F, Table S1). Our results showed that AMP drug-lock complexes are inert and lack cytotoxic activity until activation by protease activity.

### Kinetic model of prodrug treatment reveals a binary outcome

To quantitatively understand how the rate of prodrug activation competes with the number of living bacteria, we built a mathematical model using a system of nonlinear ordinary differential equations (ODE). In our system, the three dynamic populations are the Bacteria, *B*, the Locked drug, *L*, and the Unlocked drug, *U*, for which we formulated governing ODEs by considering the system parameters that affect populational change over time. For this prodrug system, increased *E. coli* growth results in increased proteolytic activation of AMP, which in turn lyses bacteria to create a negative feedback loop. Therefore, we modeled the bacteria population *B* as increasing (i.e., green dashed arrows) the rate of enzymatic drug conversion (i.e., *L* to *U*) and the unlocked drug population *U* as increasing the rate of bacterial death (Fig 2A). To account for the fact that bacterial growth rate, *r*, slows down as environmental resources become limiting (i.e., carrying capacity, *B*_*max*_), we used a logistic growth model ^35^, which produces an S-shaped curve and has been used extensively in biology to study population expansion^36^ and tumor growth^37^. By contrast, the rate of *E. coli* death is proportional to the concentration of unlocked drug, *U*, and the concentration of Bacteria, *B*, according to a proportionality constant, *a*, that represents the amount of AMP required to kill *E. coli* per unit time (Table S2, Equation 1.1).

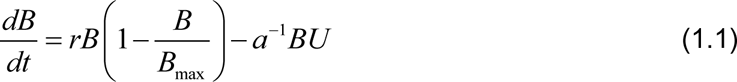

**Figure 2.**
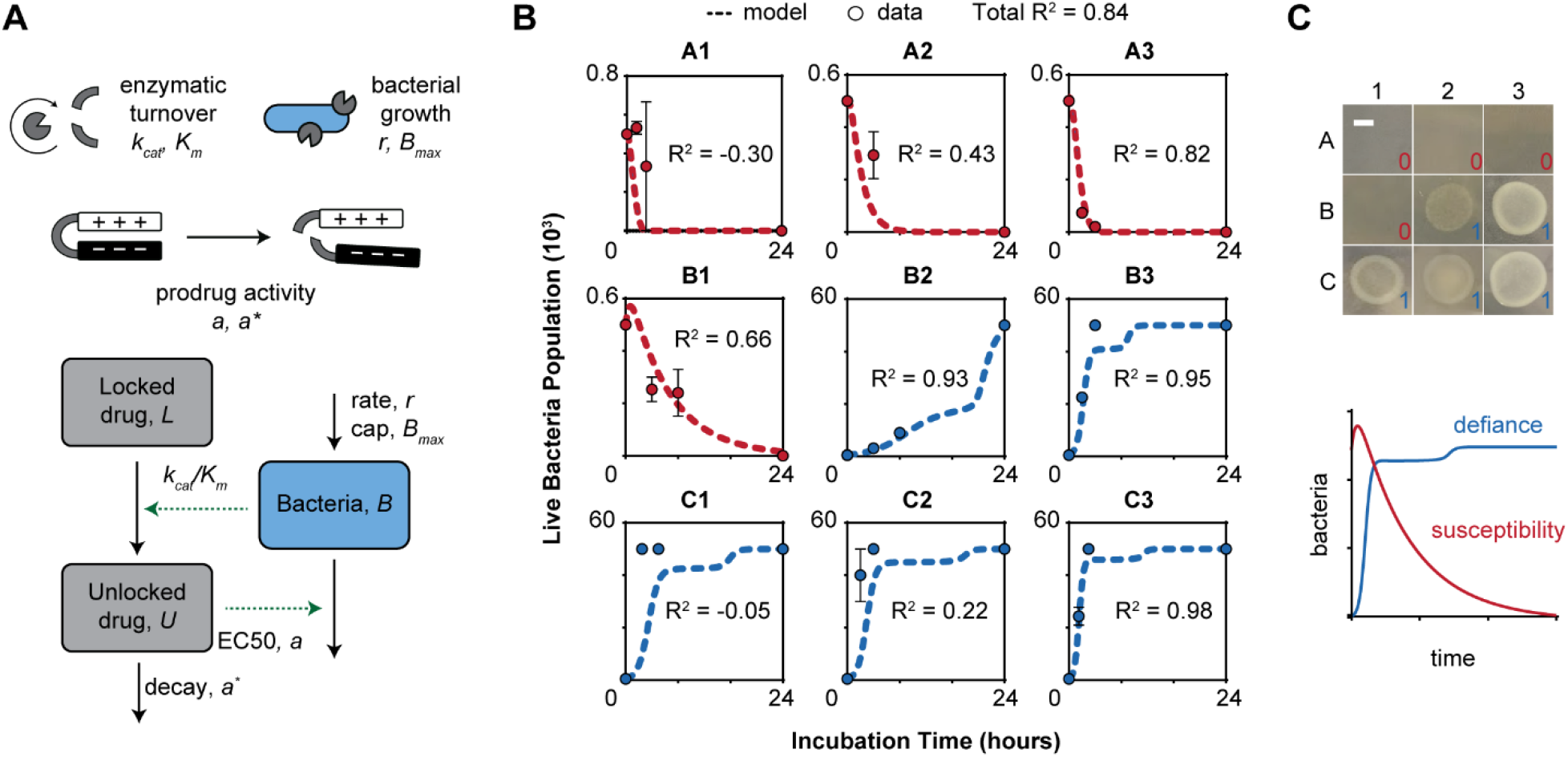
Kinetic model of prodrug defiance reveals a binary outcome. (**A**) Graphical representation of the multi-rate model of bacteria (*B*), locked drug (*L*), and unlocked (*U*) drug populations. Legend describes the agents associated with each variable. (**B**) Validating the model with serial CFU measurements (red and blue dots; n = 3, error bars SEM) and ODE model simulations of nine conditions (A1–3, B1–3, C1–3) given extracted growth rate and enzyme kinetics parameter values. (**C**) Agar plates taken at end-point showing the binary presence (1) or absence (0) of bacterial growth for nine environmental conditions (A1–C3, scale bar = 4 mm). Schematic depicting that bacterial defiance is characterized by proliferation in the presence of activated drug.

To model the rate of enzymatic activation of locked drugs, *L*, by OmpT, we applied Michaelis-Menten (MM) kinetics ^38^, where the rate of substrate activation is determined by the catalytic rate of the reaction, *k*_*cat*_, and the half-maximal substrate concentration, *K*_*m*_ (Equation 1.2). Here we assumed our system constituted a well-mixed solution of freely diffusing substrates (i.e., locked drug) in large excess, which were valid assumptions for this study since AMP prodrug was present at concentrations ∼10^2^ micromolar in an aqueous environment. Because the total amount of drug is conserved, we defined the MM activation rate of unlocked drug, *U*, as opposite of the degradation rate of locked drug *L* (Equation 1.3). We further included a term to account for the loss of active AMPs according to a proportionality constant, *a*^*\**^, that represents the number AMP required to kill one *E. coli.* This term was necessary as AMPs lyse *E. coli* by intercalating with bacterial membranes^39^ and therefore are removed from the system and unable to target additional bacteria.

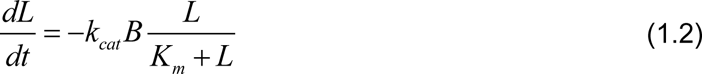

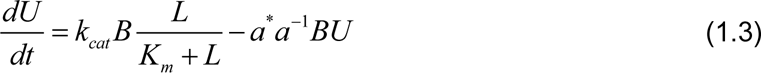

To fit this model to our system, we measured the values for relevant parameters, including enzymatic efficiency (e.g., *k*_*cat*_, *K*_*m*_), bacterial growth (e.g., *r*, *B*_*max*_), and prodrug activity (e.g., *a*, *a*^*^) (Fig. S1, Table S2). This allowed us to predict bacteria-prodrug response curves (red and blue dashed lines; Fig. 2B) across nine distinct combinations of *k*_*cat*_ and *r* values, which we experimentally controlled by altering the ambient temperature and concentration of broth (conditions labeled A1–3, B1–3, and C1–3; Table S3). Strikingly, our model anticipated a binary bacterial response to prodrug treatment that would be evident within 24 hours; bacteria were predicted to be either susceptible to the prodrug and die (red dashed lines) or to exhibit a drug-invariant phenotype and proliferate to saturating levels (blue dashed lines) (Fig. 2B, C). To experimentally validate our model predictions, we incubated bacteria with prodrugs under the defined nine conditions, and quantified the number of living bacteria longitudinally over the course of a 24 hour treatment window. Quantified bacterial counts taken during the course of treatment as well as at endpoint closely matched the values predicted by our model (R^2^ = 0.84, red and blue dots; Fig. 2B, C) and likewise revealed a binary bacterial response to prodrugs leading to antibiotic failure. We therefore called this form of resistance to prodrugs as bacterial “defiance.” Collectively, these experiments demonstrate that when *E. coli* are exposed to the AMP prodrug, our model can be used to predict bacterial growth kinetics, which ultimately result in a binary outcome.

### Predicting bacterial defiance with a dimensionless parameter

Having demonstrated that bacteria exposed to a prodrug exhibit a binary outcome – susceptibility (i.e., death) or defiance (i.e., survival) – we next sought to determine whether the behavior could be quantified by a metric of resistance that could be generalized across broad treatment conditions. Based on our model and experimental validation, we observed that under defiance, populations of live bacteria expanded (i.e., positive growth rate) throughout the course of treatment, which implied that key bacterial growth parameters (e.g., *r*, *B*_*max*_) were greater in value than enzyme-driven death rate parameters (e.g., *k*_*cat*_, *K*_*m*_) (Fig. 2B, 3A). Therefore, to derive a metric to discriminate between defiant and susceptible bacteria, we used Buckingham Pi theorem to identify a dimensionless quantity that represents the competing balance between growth rate and enzymatic turnover, which we defined as the Bacterial Advantage Heuristic (*B.A.H.*) (Equation 1.4).

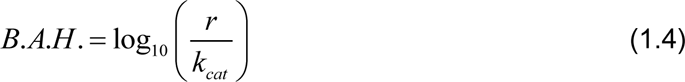

**Figure 3.**
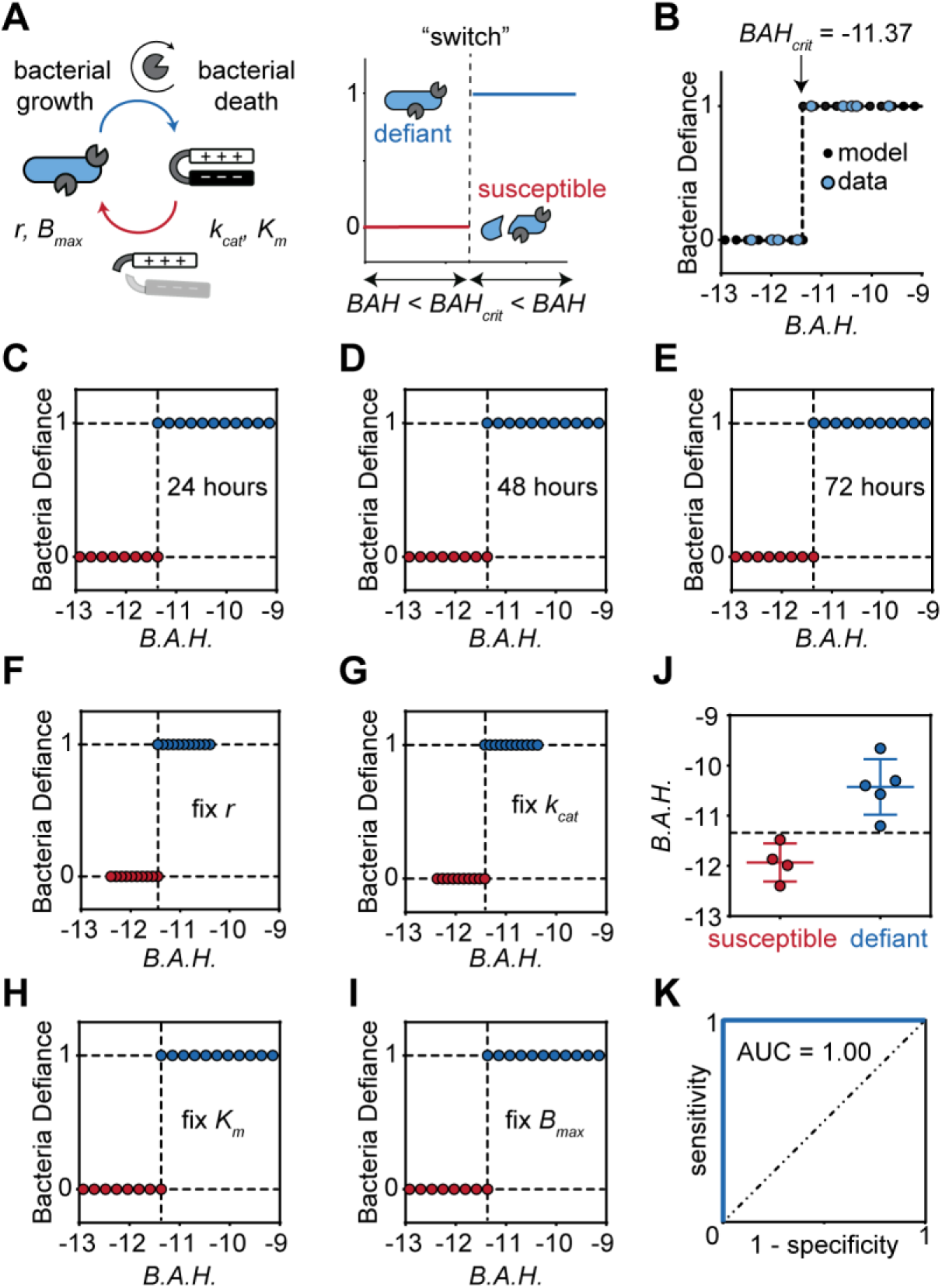
Predicting bacterial defiance with a dimensionless constant. (**A**) Schematic depicting competing rates between bacteria growth, prodrug activation, and bacteria death with the relevant parameters. Illustrative graph showing the “switch” like behavior that occurs at a critical *BAH* (*BAH*_*crit*_) threshold (bacterial defiance = 1 when BAH > *BAH*_*crit*_, susceptibility = 0 when BAH < *BAH*_*crit*_) (**B**) Model prediction (dashed line) and experimental validation (blue dots) of bacteria outcome as a function of *BAH* to validate the critical defiance value *BAH*_*crit*_. (**C–E**) Model simulations of bacteria outcome as a function of *BAH* for treatment durations 24, 48, and 72 hours. (**F–I**) Model simulations of bacteria outcome versus *BAH* by controlling four parameters: *r*, *k*_*cat*_, *K*_*m*_, and *B*_*max*_. In each panel, one of these four variables is fixed while the others are changing. (**J**) Classifying bacteria into defiant and susceptible populations based on *BAH*. (**K**) Receiver-operating characteristic (ROC) analysis using *BAH* to classify populations as defiant based on bacteria-driven parameters (*r* and *k*_*cat*_/*K*_*m*_).

This quantity reflects that bacteria have a higher probability of switching to the defiance phenotype under conditions that promote a faster growth rate, *r*, and slower enzymatic activity, *k*_*cat*_. To predict the onset of the defiance phenotype, we sought to determine the critical *B.A.H.* value that distinguishes defiant bacteria from susceptible populations. Using our mathematical model, we simulated >2000 prodrug treatment conditions covering a range of values for *r*, *B*_*max*_, *k*_*cat*_, and *K*_*m*_ and identified a critical value of BAH at which bacteria switch from susceptibility to defiance (*BAH*_*crit*_ ∼ −11.37) (Fig. 3B). In other words, for any environmental or genetic condition that produces a *BAH* > *BAH*_*crit*_, the bacteria will survive the prodrug treatment, and for *BAH* < *BAH*_*crit*_, the bacteria will die. To test the robustness of this switch-like behavior, we modeled treatment outcomes by varying bacterial growth rates (*r*) and enzymatic efficiencies (*k*_*cat*_) (> 100 points) and found that *BAH*_*crit*_ was time-independent across a range of prodrug treatment durations. (Fig 3C–E). Additionally, to demonstrate that *BAH*_*crit*_ is independent from any system parameter, we modeled treatment outcomes (> 500 points) by individually fixing each of the four parameters (i.e., *r*, *B*_*max*_, *k*_*cat*_, and *K*_*m*_) in turn and found that *BAH*_*crit*_ was invariant across all conditions tested (Fig 3F–I). To confirm the value of *BAH*_*crit*_ experimentally, we first used the nine experimental conditions previously tested (A1–3, B1–3, C1–3) to fit the values for *k*_*cat*_ and *r* (Fig S2), and then used the computationally-derived critical *BAH* threshold to classify the phenotype of nine bacteria-prodrug treatment conditions. By receiver-operating-characteristic (ROC) analysis, *BAH*_*crit*_ perfectly predicted the onset of defiance (Fig. 3J,K AUROC = 1.00, n = 9) with 100% specificity and sensitivity. Our model results predicted that changing key system parameters to decrease the *B.A.H.* below the critical threshold will result in successful treatment of defiant bacteria. To demonstrate this, we took three different AMP prodrugs with distinct linker sequences (Table S1) with increasing *k*_*cat*_ values for OmpT to decrease the *B.A.H.* below *BAH*_*crit*_, which allowed us to successfully treat previously defiant populations of bacteria (Fig. S3). These findings are important for the successful design and administration of prodrug therapies, which may be improved by optimizing fundamental pharmacokinetic parameters.

### Combating bacterial defiance with prodrug biological circuits

Our model revealed that bacterial defiance arises during prodrug monotherapy when environmental parameters, as represented by a dimensionless constant, crosses a critical transition *BAH* value. To combat bacterial defiance, we therefore sought to design a multi-prodrug approach to eliminate defiant bacteria that would otherwise resist treatment to a single prodrug (Fig. 4A). To do this, we recognized that the bacterial switch from susceptibility to defiance could be considered as behaving analogously to electronic transistors – transistors allow input current (I_in_) to pass (I_out_) when the gate voltage (G) crosses a threshold, whereas with AMP prodrugs, input bacteria (B_in_) survive treatment (B_out_) when the *BAH* (gate) crosses a critical value (Fig. 4B). Under this analogy, we postulated that multiple prodrug biological transistors could be used to construct logical operations (i.e., AND, OR, NOT gates) that allow for the design of integrated biological circuits that output state “0” (i.e., bacterial death) for all possible inputs. These biological circuits would then be representative of a multi-prodrug strategy to eliminate bacteria even when input *BAH* variables (i.e., temperature, nutrient level, enzyme activity) would result in a state of defiance for a single prodrug. To accomplish this, we designed complementary N-type and P-type transistors, which are required to construct all possible logic gates^40^. N-type transistors allow input current to pass when the gate signal is above a defined threshold, whereas P-type transistors allow current to pass when the gate signal is below a defined threshold. We constructed N-type transistors with our AMP prodrug system, which allowed input bacteria (B_in_ = 1) to survive (B_out_ = 1) when the *BAH* is above *BAH*_*crit*_ which we used to define the gate threshold (i.e., *BAH* > *BAH*_*crit*_ equivalent to G = 1; *BAH* < *BAH*_*crit*_ equivalent to G = 0) (Fig. 4B). To create P-type transistors, we synthesized heat-triggered liposomes^41^ (Fig. S4) loaded with the antibiotic ampicillin^42^, which was released at temperatures above 37°C to kill bacteria (i.e., B_out_ = 0) (Fig. 4C). Above this critical temperature, treatment with AMP prodrugs leads to bacterial defiance (i.e., BAH > *BAH*_*crit*_ or G = 1) and *E. coli* survive prodrug treatment by proliferating significantly faster than prodrug activation^43^. We therefore defined temperatures above 37 °C as corresponding to *G* = 1 and below as *G* = 0 for our P-type biological transistor to match inverse gate values for our N-type biological transistor. These results demonstrated that antimicrobial drugs could be modeled with similar input and output characteristics compared to N-type and P-type transistors.

**Figure 4.**
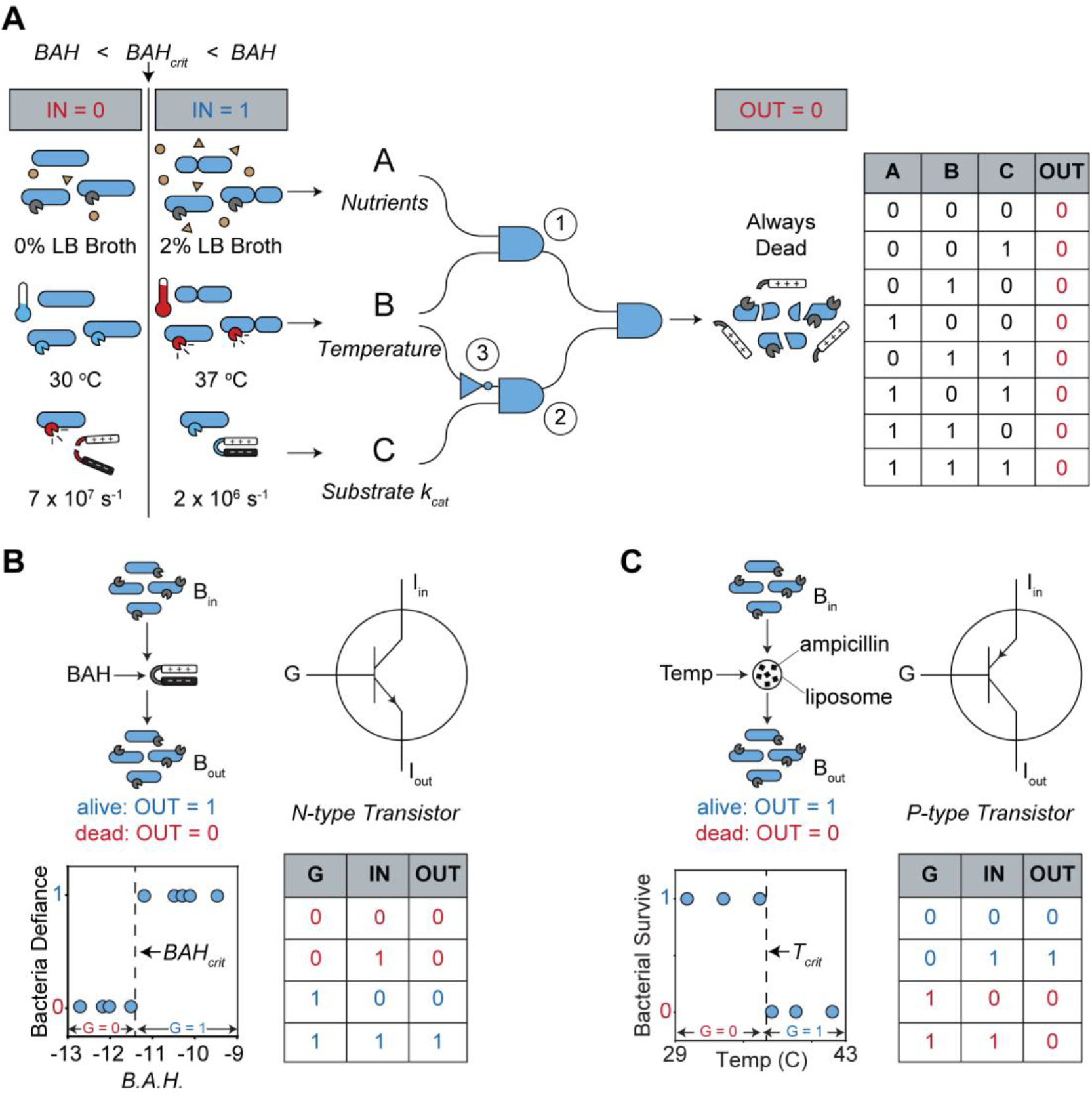
Building biological transistors with prodrugs. (**A**) Logic circuit schematic depicting a function that universally outputs 0 for all possible combinations of three binary inputs. Inputs are controlled by environmental variables including bacterial nutrients (brown circles and triangles), ambient temperature (thermometers), and substrate specificity (red substrate = high specificity, blue substrate = low specificity). Depicted parameter values represent experimental conditions. Truth table for universal circuit input-output combinations. (**B**) Schematic and truth table showing how the AMP prodrug behaves analogously to N-type transistors. Electronic N-type transistors allow the input current (I_in_) to pass when the gate voltage (G) is *high*. Similarly, the N-type prodrug transistor allows the input bacteria (B_in_) to survive when the gate signal (G) is *high* (i.e., G = 1 when *BAH* > *BAH*_*crit*_). (**C**) Schematic and truth table showing how the heat-triggered, drug loaded (black squares) liposomes (black circle) behave analogously to P-type transistors. Electronic P-type transistors allow the input current (I_in_) to pass when the gate voltage (G) is *low*. Similarly, heat-triggered, drug-loaded liposomes allow the input bacteria (B_in_) to survive (B_out_ = 1) when the gate signal (G) is *low* (i.e., G = 0 when temperature < *T*_*crit*_). All truth tables depict binary input-output combinations.

To demonstrate the use of multiple prodrug transistors in biological circuits capable of killing defiant bacteria, we selected three variables (i.e., bacterial nutrients, temperature, linker substrate *k*_*cat*_) that independently influence bacteria-prodrug competition to act as circuit inputs (i.e., A, B, and C respectively). For each circuit input, we assigned the value 0 or 1 by fixing all other variables and choosing one condition that enabled bacterial survival (i.e., IN = 1 when *BAH* > *BAH*_*crit*_) and one condition that resulted in bacterial death (i.e., IN = 0 when *BAH* < *BAH*_*crit*_). We then designed a biological circuit to kill bacteria (i.e., OUT = 0) under all eight combinations of nutrient (LB Broth = 0% or 2%), temperature (T = 30 °C or 37 °C), and linker substrate (*k*_*cat*_ = 7 × 10^7^ s^−1^ or 2 × 10^6^ s^−1^) conditions while using three AND gates and one NOT gate (Fig 4A). Electronic AND gates are created by placing two transistors in series; similarly, we sequentially dosed bacteria with two prodrugs, each with corresponding *BAH* values to represent the gate inputs (i.e., *A* and *B*). We controlled the corresponding *BAH* value for each prodrug with the linker substrate sequence (e.g., A or B = 0 when *k*_*cat*_ = 7 × 10^7^ s^−1^, A or B = 1 when *k*_*cat*_ = 2 × 10^6^ s^−1^) and tested all four combinations of the two prodrugs (i.e., AB = 00, 01, 10, and 11). When exposed to all four possible inputs, bacteria only survived the condition with two low *k*_*cat*_ prodrugs (i.e., AB = 11), which matched the ideal outputs of an AND gate (Fig. 5B). These results show that dosing multiple prodrugs in sequence reduces the fraction of bacterial populations that survive (e.g., one prodrug = 50% survival, two prodrugs = 25% survival, etc.). Whereas the AND gate comprised prodrugs in series, we demonstrated one implementation of an OR gate by splitting a population of bacteria in half (i.e., separate wells), dosing each with a different prodrug during the same time interval (i.e., in parallel) and recombining the bacteria post-treatment. Using this circuit, bacteria survived in any case where at least one half of the population was dosed with low *k*_*cat*_ prodrug (i.e., AB = 01, 10, or 11), matching the ideal outputs of an OR gate (Fig. 5C). We created a NOT gate by using a P-type transistor comprising ampicillin-loaded heat-triggered liposomes, which caused bacteria to die at high temperatures (i.e., IN = 1) and survive at low temperatures (i.e., IN = 0). (Fig. 5D). These results show that heat-triggered liposomes can be used as a fail-safe to kill bacteria above the critical temperature representing the onset of the defiance phenotype. To validate the multi-prodrug circuit, we incubated populations of bacteria under each of the eight environmental conditions with three prodrugs: (1) an AMP prodrug with fixed *k*_*cat*_/*K*_*m*_, (2) an AMP prodrug with *k*_*cat*_/*K*_*m*_ determined by input C, and (3) a heat-triggered, drug-loaded liposome (Fig. 4A). Whereas a single prodrug only eliminated bacteria in half of the input cases (Fig. 5E), our circuit autonomously eliminated bacteria under all eight combinations of the three environmental signals (Fig. 5F). These results demonstrate that by utilizing the transistor-like properties of prodrugs, we can design and administer biocircuits that combat bacterial defiance.

**Figure 5.**
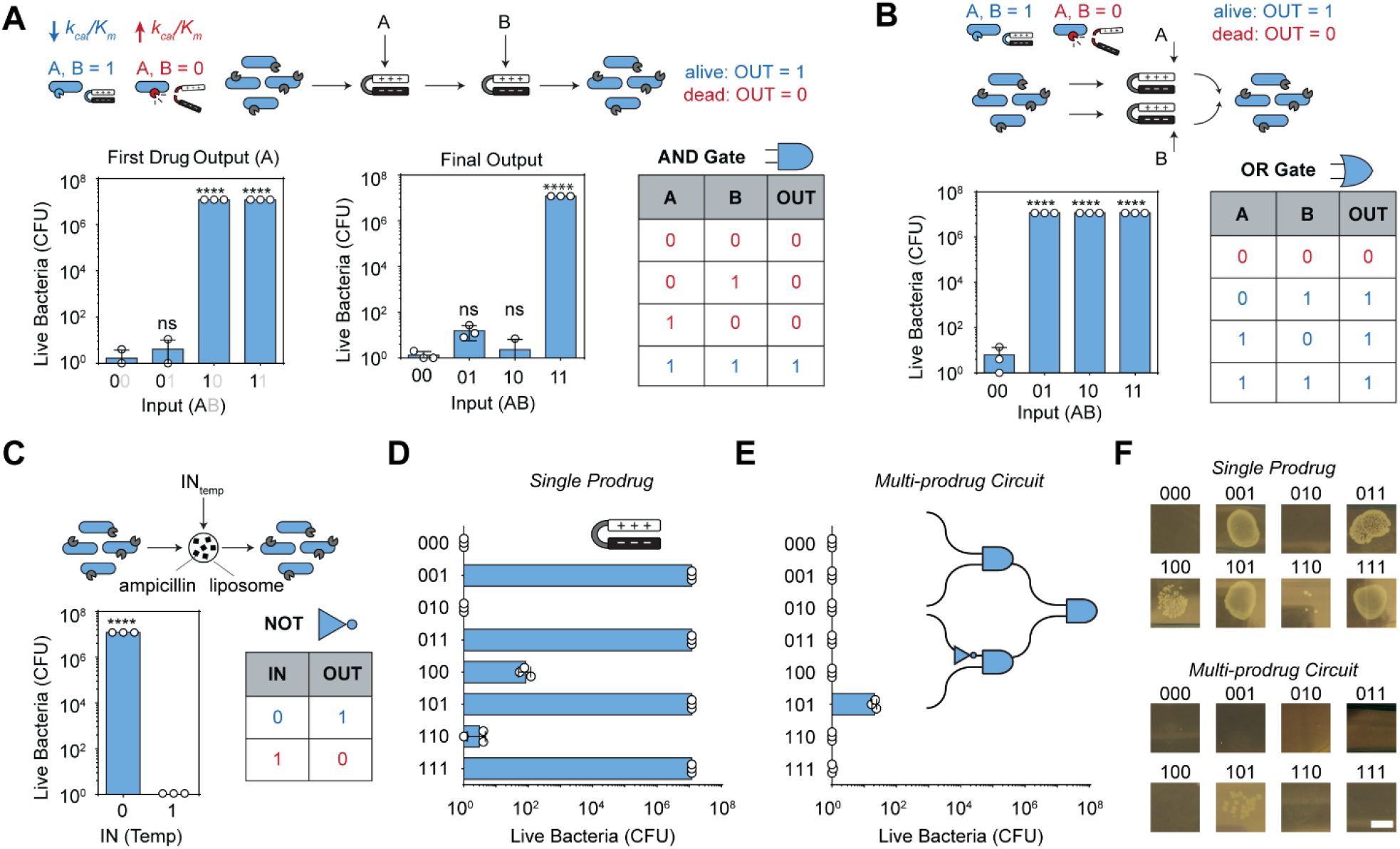
Combating bacterial defiance with a logic-based prodrug circuit. (**A**) Bacterial viability assay validating the function of a prodrug-based AND gate. Schematic depicts two prodrug transistors arranged in series. Blue substrate represents low preference (*k*_*cat*_) for bacterial protease OmpT (blue = low activity), and red substrate represents high preference (*k*_*cat*_) for bacterial protease OmpT (red = high activity). (**B**) Bacterial viability assay validating the function of a prodrug-based OR gate. Schematic depicts two prodrug transistors arranged in parallel. For AND and OR, input values were controlled with substrate sequence, where the substrate with high k_cat_/K_m_ corresponds to *BAH* = 0 and the substrate with low K_cat_/K_m_ corresponds to *BAH* = 1. Statistical comparisons are made with 00 condition. (**C**) Bacterial viability assay validating the function of a prodrug-based NOT gate. Schematic depicts heat-triggered liposome (black circle) loaded with drug (black squares = ampicillin). BAH = 0 corresponds to temperature of 30 °C and BAH = 1 corresponds to 37 °C. All truth tables depict ideal binary input-output combinations. (**D**) Bacterial viability assay quantifying the outcome of a single prodrug in the context of different environmental conditions. (**E**) Bacterial viability assay demonstrating the eight outputs of the autonomous circuit. (**F**) Representative CFU images for single prodrug and multi-prodrug treatment (scale bar = 4 mm). Error bars represent standard deviation (n = 3). * < 0.05, ** < 0.01, *** < 0.001, and **** < 0.0001.

## DISCUSSION

The rising threat of antibiotic failure has motivated the classification of resistance mechanisms to improve the design of future treatment strategies. In this work, we identified a prodrug-specific subclass of resistance, which we named defiance. To study bacterial defiance, we synthesized an AMP prodrug that was activated by bacterial protease activity and successfully killed *E. coli*. However, we found that this prodrug failed in conditions where the bacteria grew faster than the rate at which they activated the drug. We developed a mathematical model of the bacteria-prodrug competition and derived a dimensionless constant (*BAH*), which we used to perfectly predict the onset of the defiance phenotype. These studies revealed that both environmental (e.g., temperature, available nutrients) and pharmacokinetic (e.g., activation rate *k*_*cat*_) parameters can be tuned to engineer successful prodrug therapies. In future work, this information could be leveraged to improve the efficacy of existing prodrugs by tuning the prodrug decay constant (i.e., biological half-life), which directly changes the *BAH*_*crit*_ value. Alternatively, the catalytic efficiency of the prodrug substrate could be tuned to change the *BAH* value and increase the probability of success, which has been previously demonstrated by engineering prodrug substrates with higher affinity for the enzymatic target ^41, 45^.

In our study, insights into bacterial defiance were elucidated in the context of AMP prodrugs that activate by an extracellular mechanism of action. We anticipate that this phenomenon of defiance also extends to prodrugs that are intracellularly activated, which represent the second major category of prodrugs^46^. To account for intracellular activation, our model would include a transport rate term to account for prodrug diffusion across cell membranes ^47^. The implications of our model will likely still hold if prodrug transport is not the rate limiting step (i.e., represents a separate mechanism by which bacteria can escape treatment) and is greater than the rate of prodrug activation. This is a reasonable assumption given that the rates of drug diffusion across membranes ^48–49^ are on the order 10^10^–10^17^-fold higher than the typical range of prodrug activation rates ^50, 51^.

By comparison to established forms of resistance, prodrug defiance could occur from either genetic or phenotypic changes. For example, both phenotypic resistance (i.e., non-inherited resistance) and tolerance are acquired under specific environmental conditions that affect growth rate, such as in biofilms ^6, 52^. By contrast, inherited resistance arises from genetic mutations that alter key bacterial processes including metabolism, enzymatic activity, and drug efflux pumps ^53^. In our studies, we demonstrated that environmental conditions (i.e., temperature, nutrient levels) affect the defiance phenotype; however, it is also likely that the onset of defiance could also arise from genetic mutations that affect E. coli growth rates or cleavage activity or proteases^54–56^. Our findings suggest that prodrugs currently used in the clinic may fail due to the onset of bacterial defiance. For example, the nitroimidazole class of antibiotics (e.g., metronidazole, dimetridazole, tinidazole, etc.), which is used to treat anaerobic bacteria (e.g., *Enterococcus* species, *Clostridium* species, *Helicobacter pylori*, etc.) represent prodrugs that are activated by bacterial reductases ^57^. Genetic studies have revealed that bacterial resistance to nitroimidazole antibiotics is caused by either partial or complete reduction in expression of genes (e.g., rdxA, frxA, etc.) encoding the reductases that activate the prodrug ^58–60^. These changes in the activating enzyme would decrease the rate of prodrug activation, thereby resulting in a reduced overall *BAH*, that if below *BAH*_*crit*_ for the system, would result in defiance.

As bacteria-activated prodrugs are an emerging strategy to treat antibiotic resistant infections ^9^, our work may provide strategies for preventing defiance-related resistance mechanisms in new and existing prodrugs. To propose strategies for treating prodrug defiance, we leveraged the transistor-like behavior of prodrugs to design a multi-drug circuit to eliminate defiant bacteria that would normally survive the treatment of a single prodrug. In these experiments, we tested eight discrete combinations of three variables (e.g., nutrients, temperature, and linker substrate) and showed that the multi-drug logic circuit effectively eliminated bacteria in all eight conditions. In future work, different variables (e.g., O_2_, pH, relative biodistribution, etc.) could be included to increase the range of possible treatment conditions. This would require a quantitative understanding of the relationship between the variable and how it affects key components of the BAH (i.e., *r* or *k*_*cat*_). Ideally, these variables will proportionally influence either the growth rate or the enzymatic activity of the bacteria, such as how increasing substrate efficiency increases *k*_*cat*_, which creates one critical value (*BAH*_*crit*_). By contrast, variables that influence growth rate or enzyme activity with multiple critical values (i.e., inflection points) limit the range of applicability. For example, higher temperatures increase both growth rate and enzymatic activity, but above critical temperatures, protein enzymes denature, resulting in irreversible loss of activity ^61^. However, these values are beyond relevant biological values and do not limit the applicability of these prodrug treatments to normal physiological conditions.

Here, we quantitatively studied the mechanism of prodrug defiance and revealed logic-based strategies for successfully treating bacteria. We envision that this body of work will motivate the development of new prodrugs and improve the clinical administration of existing prodrugs, ultimately helping to reduce the burden of antibiotic failure.

## Author Contributions

G.A.K. and B.A.H conceived of the idea. G.A.K, B.A.H., and I.C. designed experiments and interpreted results, and wrote the manuscript. B.A.H. and I.C. carried out experiments.

## Acknowledgments

This work was funded by an NIH Director’s New Innovator Award (Award No. DP2HD091793). B.A.H is supported by the NSF GRFP, National Institutes of Health GT BioMAT Training Grant under Award Number 5T32EB006343 and the Georgia Tech President’s Fellowship. This material is based upon work supported by the National Science Foundation Graduate Research Fellowship under Grant No. DGE-1650044 (B.A.H.). G.A.K. holds a Career Award at the Scientific Interface from the Burroughs Welcome Fund. This work was performed in part at the Georgia Tech Institute for Electronics and Nanotechnology, a member of the National Nanotechnology Coordinated Infrastructure (NNCI), which is supported by the National Science Foundation (Grant ECCS-1542174). The content is solely the responsibility of the authors and does not necessarily represent the official views of the National Institutes of Health. The authors would like to thank Ian Miller (Georgia Tech), Shreyas Dahotre (Georgia Tech), Dr. David Myers (Georgia Tech & Emory), and Dr. Kyle Allison (Georgia Tech & Emory) for their helpful discussions.

## Data Availability

The authors declare that the data supporting the findings of this study are 20 available within the paper and its supplementary information files.

## Code Availability

The authors declare that the code supporting the findings of this study is available within the supplementary information files.

## EXPERIMENTAL PROCEDURES

### Protease Cleavage Assays

All protease cleavage assays were performed with a BioTek Cytation 5 Imaging Plate Reader, taking fluorescent measurements at 485/528 nm (excitation/emission) for read-outs measuring peptide substrates terminated with FITC (Fluorescein isothiocyanate). Kinetic measurements were taken every minute over the course of 60 – 120 minutes at 37 C. Tobacco Etch Virus protease (TEVp), along with its substrate and buffer was obtained from Anaspec, Inc. (Fremont, CA). Activity RFU measurements were normalized to time 0 measurement, and as such represent fold change in signal. Outer Membrane Protease T (i.e., OmpT, Protease 7) was purchased from Lifespan Biosciences (Seattle, WA). OmpT fluorescent peptide substrate was custom ordered from Genscript (Piscataway, NJ).

### Bacterial culture and cytotoxicity measurement

DH5α Escherichia coli were a gift from Todd Sulchek’s BioMEMS lab at Georgia Tech. E. coli were cultured in LB broth (Lennox) at 37 C and plated on LB agar (Lennox) plates. LB broth was purchasd from Millipore Sigma (Burlington, MA) and LB agar was purchased from Invitrogen (Carlsbad, CA). AMP and locked AMP were custom ordered from Genscript (Piscataway, NJ). See Table S1 for more information. Bacteria were grown to a concentration of 10^9^ CFU/mL before being used for experiments. Concentration was estimated by measuring the OD_600_ of the bacterial suspension, and assuming an OD_600_ of 1.000 corresponds to a concentration of 8 × 10^8^ CFU/mL. Bacterial cell viability was measured by making eight 10-fold serial dilutions, and plating three 10 uL spots on an LB agar plate. Plates were incubated overnight at 37C, and CFUs were counted. Untreated bacteria CFU counts served as control for 0% cytotoxicity, and bacteria + IPA (or 0 countable CFUs) served as control for 100% cytotoxicity.

### Liposome Preparation

DPPC and MPPC were purchased from Avanti Polar Lipics, inc. The heat-triggered NOT gate formulation was composed of DPPC:MPPC at the molar ratio 9:1, and were evaporated in cholorform to produce a lipid film. Films were rehydrated in 100% water containing 100 mg/mL ampicillin and were sonicated for 20 minutes. Lipid formulations were then purified using a PD-10 desalting column (GE Healthcare) and size was measured via dynamic light scattering (DLS, Malvern).

### Computational model

The ODE modelling and solutions were performed in MATLAB 2016b. Code can be found in supplementary information.

### Statistical Analysis

Statistical analysis was performed using statistical packages included in GraphPad Prism 6. To assess the significance of increase in signal due to protease cleavage, we used a two-way ANOVA (without repeated measures) followed by Sidak’s multiple comparisons test. A one-way ANOVA followed by Dunnett’s multiple comparisons test was used to compare experimental means to cells only control bacterial viability assays. Two-way ANOVA followed by Sidak’s multiple comparisons test used to compare experimental means to control for bacterial cytotoxicity at multiple starting concentrations.

## SUPPLEMENTARY INFORMATION

### Measuring kinetic parameters for the ODE model

To measure drug cytotoxicity, we quantified the percent of viable bacteria when exposed to unlocked AMP drug doses ranging from nanomolar to millimolar concentrations and observed a spike in cytotoxicity in the micromolar range (Fig. S1A). To calculate bacterial growth rate constants, we incubated bacteria in broth or water at 37 °C and measured OD_600_ over the course of 8 hours, revealing a higher growth rate in broth consistent with published values^35, 56^ (3.0 > 0.1 h^−1^) (Fig. S1B). To quantify enzymatic velocity, we measured proteolytic velocity (moles L^−1^ s^−1^) under substrate concentrations ranging from 0—200 uM and fit the Michaelis-Menten equations to this data to calculate *k*_*cat*_ and *K*_*m*_ (Fig. S1C). Using these measured parameters (Table S2), we generated kinetic curves for bacteria number and drug concentration over time. The model showed that after a period of initial bacterial growth, the lagging drug population eliminated the infection burden. While the initial reservoir of locked drug concentration was ∼200 µM, the unlocked drug never rose above 5 µM (Fig. S1D), revealing that this system maintains the minimal drug concentration required to kill the infection.

**Fig. S1.**
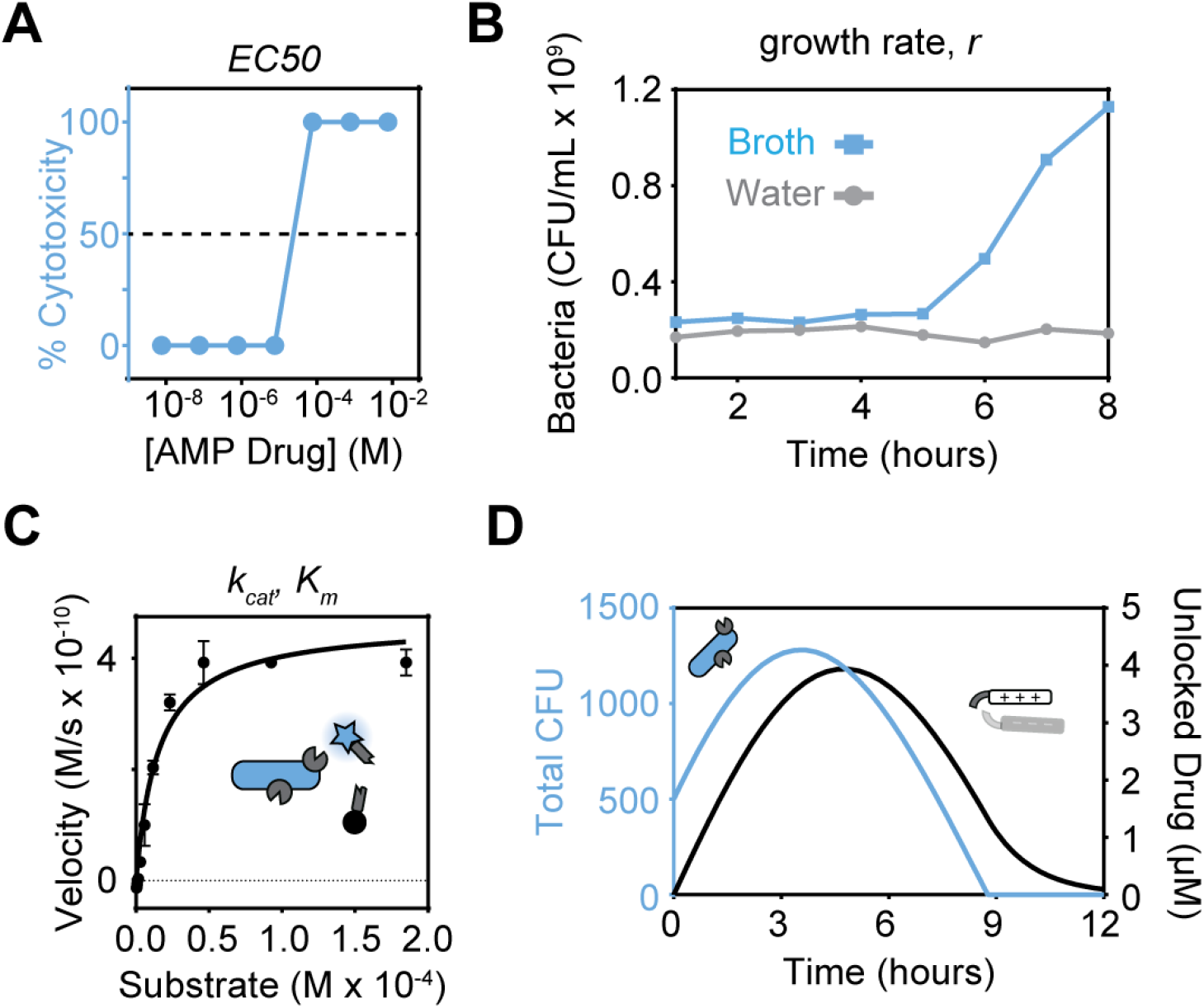
Measuring kinetic parameters for the ODE model. (**A**) EC50 measurement on a sample of *E. coli* exposed to increasing concentrations of AMP. (**B**) Growth rate calculation of *E. Coli* in broth and water by measuring OD at 600 nm. (**C**) Measuring k_cat_/K_m_ from a series of cleavage assays incubating *E. coli* at a range of substrate concentrations. (**D**) Predicting the dynamics of the bacteria and unlocked drug population given empirical parameter values. Error bars are standard deviation (n = 3).

### Estimating bacterial growth rate and enzymatic activity with parameter fitting

To estimate bacterial growth rate (*r*) and enzymatic activity (*k*_*cat*_), we used parameter fitting methods to a simplified model without a prodrug (Fig. S3). In this model, the prodrug is substituted with a fluorescent probe (*P*), which is activated with the same substrate sequence as the AMP prodrug. To simplify the model, we defined one bacterium, with ∼ 2.5 × 10^3^ OmpT proteases per cell ^62^, as equivalent to one effective enzyme. The concentration of cleaved fluorescent probe is monitored over time, and the bacterial concentration is quantified at serial time points. We fit this data to the two equations modelling bacterial growth and activation of the fluorescent probe (Equations S1.1, S1.2).

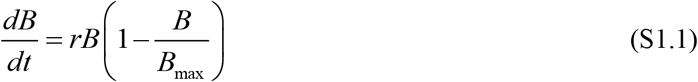

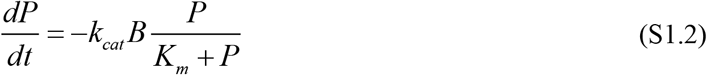

**Fig. S2.**
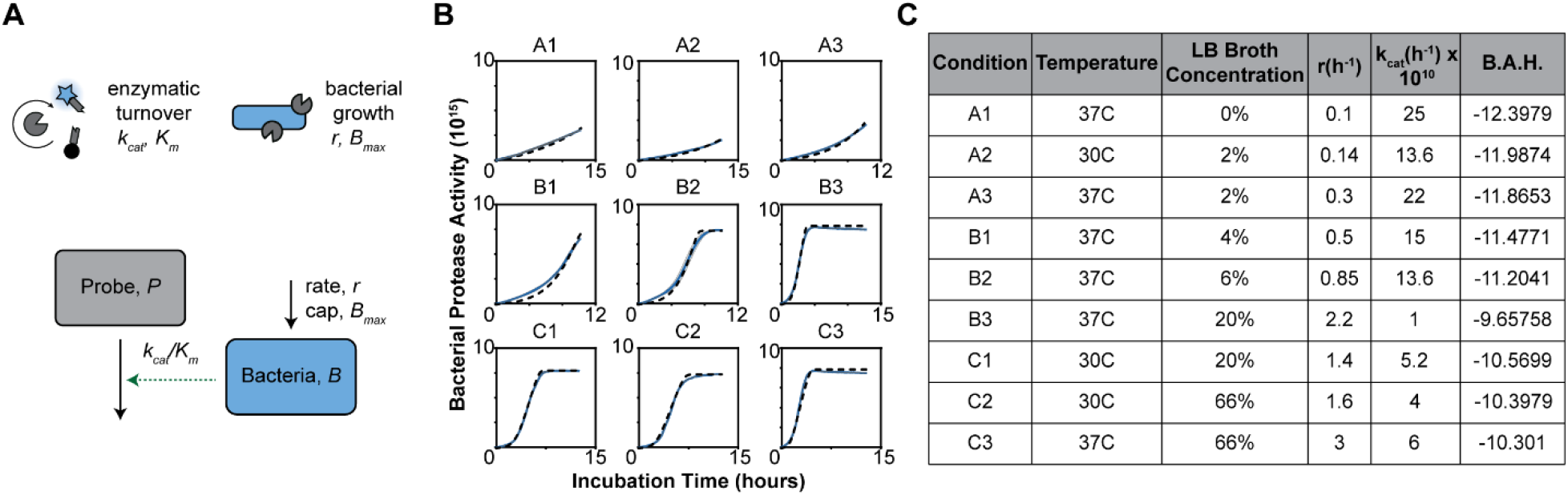
Estimating bacterial growth rate and enzymatic activity with parameter fitting. (**A**) Schematic of model used to quantify <u>*r* and</u> *k*_*cat*_ from bacterial cleavage assays. (**B**) Bacterial cleavage assays (blue line), from which growth rate (r) and enzyme kinetics (k_cat_/K_m_) are extracted, and *BAH* is calculated (inset). Dotted line represents model after parameter fitting. (**C**) Table of *r* <u>and</u> *k*_*cat*_ values quantified from model in each condition with varying temperature and LB broth concentration.

**Fig. S3.**
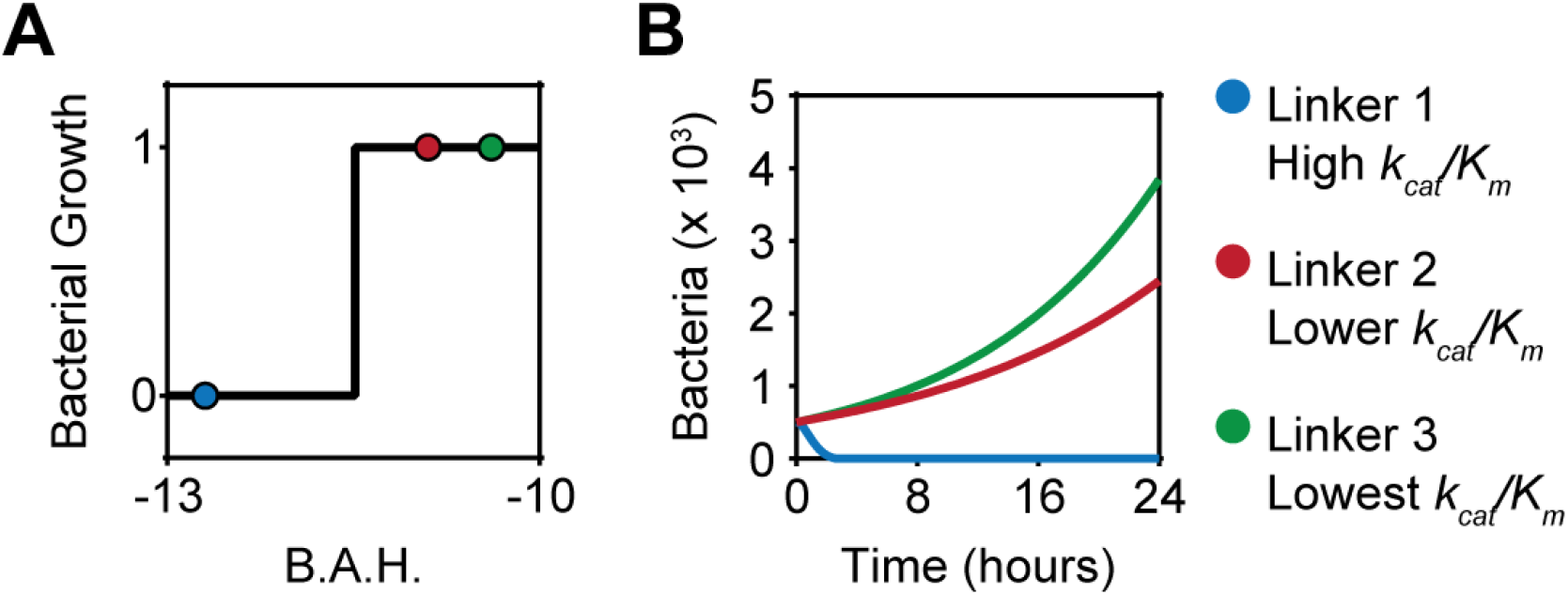
Altering drug efficacy with substrate affinity. (**A**) Bacteria viability assay post 24 hour incubation with drug unlocked by various substrates. Bacterial growth = 1 if colonies were present after plating, and bacterial growth = 0 if there were no colonies present (n = 3). B.A.H. values extrapolated from bacteria + probe velocity measurements fitted to model. (**B**) Bacteria dynamics predicted by model given three different enzyme kinetics. Linker 1 = RRSRRV, Linker 2 = RKTR, Linker 3 = ENLYFQG.

**Fig. S4.**
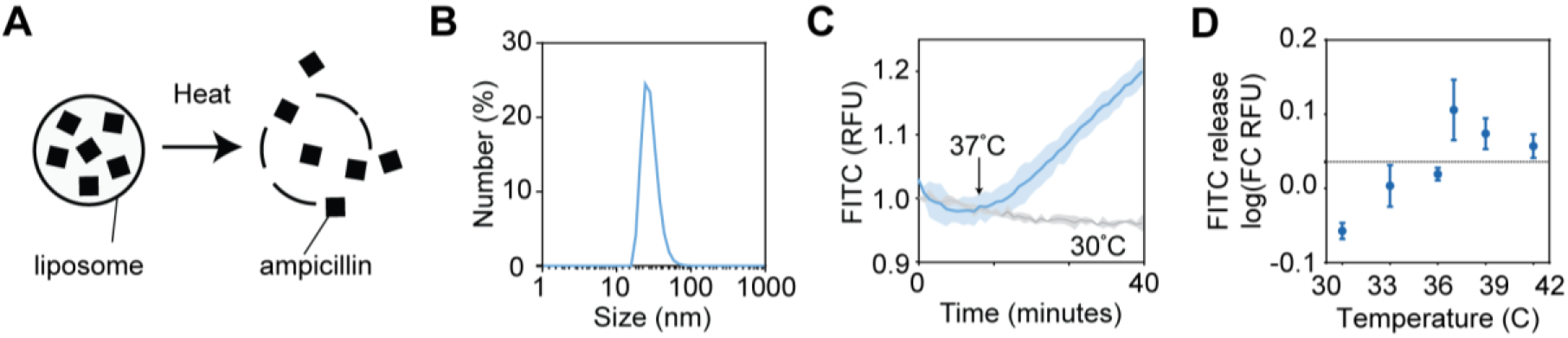
Validating heat-triggered liposomes. (**A**) Schematic depicting function of heat-triggered, drug loaded liposomes. Circles represent liposomes, black squares represent drug (ampicillin). (**B**) Dynamic light scattering of liposome formulation to determine particle size (∼35 nm). Histogram is plotted as the mean of independent measurements (n = 3). (**C**) Fluorescent assay measuring the heat-triggered release of liposome contents. Liposomes were loaded with 100 mM FITC, and heated from 25 °C to 37 °C (indicated by arrow). Grey line represents 30 °C control. Lines are plotted as means of independent measurements, shading represents standard deviation (n = 3). (**D**) Fluorescent assay measuring the release of FITC from heat triggered liposomes at various temperatures. All samples were heated to 30 °C for 10 minutes, then heated at the plotted temperature for 30 minutes. Measurements are plotted as fold change in RFU between final measurement and measurement at 30 °C. Error bars represent standard deviation (n = 3).

**TABLE S1.**

| Name | Peptide Sequence |
| --- | --- |
| Locked AMP (linker 1) | EEEEEEEEEEEEERRSRRVRRRRRRRRRR |
| Locked AMP (linker 2) | EEEEEEEEEEERKTRRRRRRRRRR |
| Locked AMP (linker 3) | EEEEEEEEENLYFQGRRRRRRRRRR |
| Locked AMP Probe | DABCYL-EEEEEEEEEEEEERRSRRVRRRRRRRRRR[Lys(5-FAM)] |
| OmpT Probe | DABCYL-RRSRRV-Lys(5-FAM) |

**TABLE S2.**
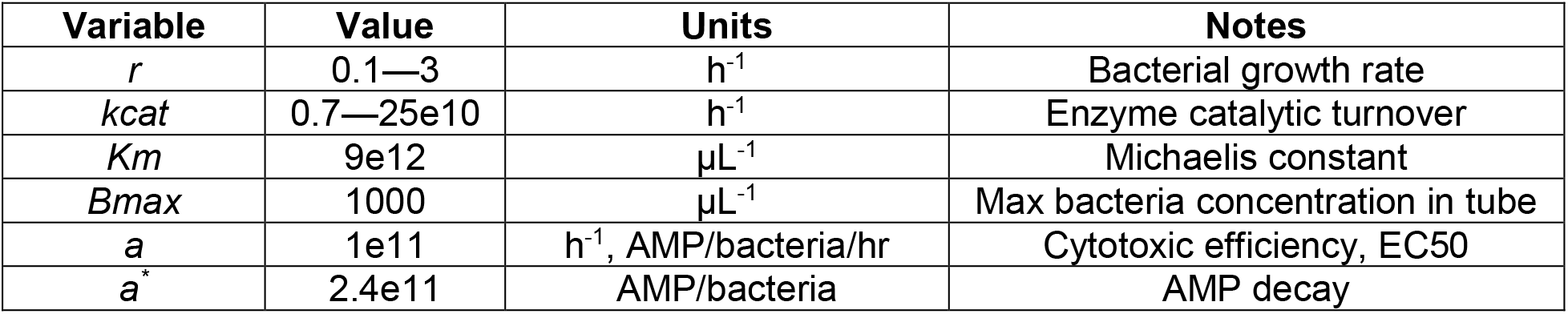

| Variable | Value | Units | Notes |
| --- | --- | --- | --- |
| $r$ | 0.1—3 | $\text{h}^{-1}$ | Bacterial growth rate |
| $k_{cat}$ | 0.7—25e10 | $\text{h}^{-1}$ | Enzyme catalytic turnover |
| $K_m$ | 9e12 | $\mu\text{L}^{-1}$ | Michaelis constant |
| $B_{max}$ | 1000 | $\mu\text{L}^{-1}$ | Max bacteria concentration in tube |
| $a$ | 1e11 | $\text{h}^{-1}$, AMP/bacteria/hr | Cytotoxic efficiency, EC50 |
| $a^*$ | 2.4e11 | AMP/bacteria | AMP decay |

**TABLE S3.**

| Location (row, column) | Temperature | LB Broth Concentration | $r (\text{h}^{-1})$ | $k_{cat} (\text{h}^{-1}) \times 10^{10}$ |
| --- | --- | --- | --- | --- |
| A1 | 37C | 0% | 0.1 | 25 |
| A2 | 30C | 2% | 0.14 | 13.6 |
| A3 | 37C | 2% | 0.3 | 22 |
| B1 | 37C | 4% | 0.5 | 15 |
| B2 | 37C | 6% | 0.85 | 13.6 |
| B3 | 37C | 20% | 2.2 | 1 |
| C1 | 30C | 20% | 1.4 | 5.2 |
| C2 | 30C | 66% | 1.6 | 4 |
| C3 | 37C | 66% | 3 | 6 |

